# Inductive bias anchors a scaling law for biological learning

**DOI:** 10.64898/2026.11.08.765173

**Authors:** Ling-Qi Zhang, Nakul Verma, Luke T. Coddington, Joshua T. Dudman, Kristin Branson, Ann M. Hermundstad

**Affiliations:** Janelia Research Campus, Howard Hughes Medical Institute, Ashburn, VA, USA; Department of Computer Science, Columbia University, New York, NY, USA

## Abstract

The speed of animal learning varies substantially across individuals and task domains, and does not always track intuitive notions of task complexity. A nominally simple binary decision task can take thousands of rewards to reach proficiency, whereas seemingly complex naturalistic tasks can be learned within dozens. A long-standing explanation is that learning speed reflects the alignment between an animal’s inductive biases and the task at hand. However, without a precise notion of alignment, it is difficult to translate this idea into quantitative predictions. Here, modeling an animal as a Bayesian learner that improves through iterative reward feedback, we use statistical learning theory to derive a scaling law for biological learning. The law makes two predictions. First, inductive bias multiplicatively sets the speed of learning through a single quantity, the *effective complexity*. This quantity measures how well the task aligns with the animal’s prior, and corresponds to the number of bits required to represent the task solution under that prior. We demonstrate this multiplicative scaling in a human motor learning dataset, where behavioral measurements *before learning* can precisely predict subsequent learning speed. Second, task performance improves as a power law in experience. We validate this power law in a large-scale mouse learning dataset and show that a two-parameter rescaling collapses individual learning curves onto a universal function predicted by the theory. These results suggest that behavioral measurements before learning can be used to predict task-dependent learning speeds, and they explain why learning speed reflects alignment with prior knowledge rather than nominal task complexity.

## Introduction

Animal learning can occur at vastly different speeds [1]. Yet the ease of learning does not always align with our intuitive notion of task complexity, even in one of its most basic forms: trial and error learning through reward feedback [2]. A nominally “simple” binary decision task can take thousands of rewards over weeks for an animal to reach proficiency [3], whereas rapid learning can emerge with mere dozens of rewards in seemingly difficult, naturalistic settings with multiple uncertain options [4, 5, 6]. As a result, we lack a principled way to predict how quickly an animal will learn a given task, or to design a task that will elicit rapid learning.

A long-standing explanation for the observed variability in learning speeds is that learning is faster when the task matches what the learner already knows. For example, Dubey et al. [7] found that human players took far longer to learn an Atari-style video game after removing information tied to their prior knowledge about objects and the physical world. Similarly, rats have been shown to quickly learn to associate a taste, but not a light or sound, with subsequent illness [8], suggesting that animals are “prepared” to learn some associations more easily than others [9]. These findings highlight that humans and other animals do not learn *tabula rasa*; rather, they come with assumptions about which sensorimotor and reward contingencies are more likely to occur in their environment, shaped by their evolutionary history, ecological niche, and individual experience. However, this also exposes a fundamental trade-off known in learning theory [10]: building assumptions into a learner can accelerate learning when those assumptions are correctly matched to the task, but can slow learning when they are mismatched. But without a precise way to measure these assumptions, this trade-off is difficult to quantitatively relate to behavior.

Formally, these built-in assumptions are called *inductive biases*, a central concept in statistical learning theory [11, 12]. Early work used this concept to explain how overparameterized models are able to generalize to new data [13]. More recently, this concept has been used to explain why the performance of deep learning models improves as a power law with the amount of data, model size, and compute, phenomena known as scaling laws [14, 15, 16]. One influential line of work formalizes inductive bias as the alignment between tasks and the high-variance directions in the representations of neural networks or kernel machines [16, 17]. While these theories have proven useful for understanding the inductive bias at the level of neural representations [18, 19], they do not readily translate to behavior.

A more natural formalization for behavior comes from Bayesian models of cognition [20], which treat inductive bias as a prior distribution over hypotheses that the learner brings to the task. This approach has revealed, for example, that humans are biased toward simplicity in learning logical rules [21, 22] and in choosing between competing explanations [23, 24]. More broadly, this framework has been extended to explain how innate and learned knowledge support fast learning and generalization [25, 26]. However, because these models require specifying a prior and a learning rule for each task, they do not yield a general account of how inductive bias quantitatively shapes learning speed.

Here, we bring together these two traditions within the probably approximately correct (PAC)-Bayesian framework, which combines the Bayesian formalism with the guarantees of statistical learning theory [27, 28, 29]. We consider a class of contextual bandit problems that describe many trial-based laboratory tasks. We then construct a Bayesian learner that maintains an inductive bias in the form of a prior distribution over hypothesized behavioral policies. By analyzing the behavior and performance of this learner, we derive a scaling law for biological learning that holds for any prior and any task within this class. Because the theory is formulated in terms of the learner’s behavioral policy, it can be tested directly against learning behavior. Two predictions follow. First, inductive bias determines the amount of experience required to learn, and acts multiplicatively through a single quantity: the *effective complexity* of the task for the learner, which captures the degree of alignment between the learner’s inductive bias and the task. Second, task performance improves as a power law in experience, providing a theoretical perspective on the long-proposed but contested “power law of practice” [30, 31, 32]. We validate these predictions in humans learning a motor control task, and in mice learning a visually-guided decision task. These results show how inductive bias can be measured from empirical data and used to predict learning speed, and they explain why learning speed reflects task alignment rather than nominal task complexity.

## Results

### Theoretical framework for quantifying task-dependent learning dynamics

Learning is an inherently dynamic process that depends not only on the task, but also on the assumptions that the learner brings, and on the rules governing how behaviors are modified over time. Here, we develop a framework to show that under a set of precise but general conditions, we can derive quantitative predictions about how the alignment between task structure and the learner’s assumptions shapes learning dynamics driven by trial-by-trial, reward-based feedback.

Specifically, we formalize a typical laboratory learning task as a contextual bandit problem [33], and we model an animal as a Bayesian learning agent (Fig. 1A). On each trial *i* of the task, the agent observes a sensory context *s*_*i*_ ∼ *ρ*(·) sampled from an unknown distribution. The agent then takes an action *a*_*i*_ and receives a reward *r*_*i*_ = *R*(*s*_*i*_, *a*_*i*_) according to a fixed reward rule. We assume the behavior of the agent is described by a family of policies *π*_*θ*_(*a* | *s*) that map sensory contexts to actions, and that can be updated over time based on experience. To this end, the agent maintains and updates a belief *q*_*i*_(*θ*) over policies, and acts by first sampling policy parameters *θ*_*i*_ ∼ *q*_*i*_(·), and then sampling an action 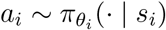.

**Figure 1:**
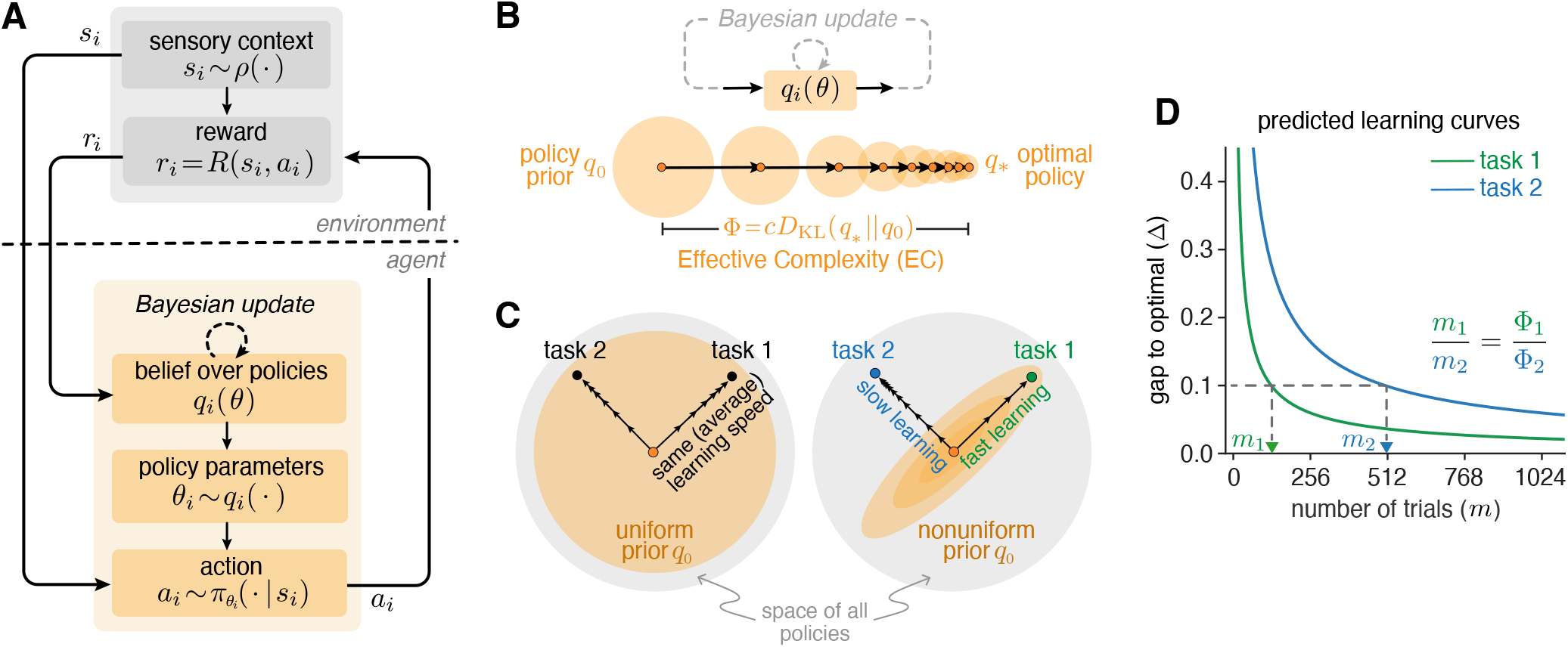
A scaling law derived from a theoretical analysis of Bayesian learners in contextual bandit tasks. **A)** A typical laboratory task modeled as a contextual bandit. On each trial *i*, the environment presents a sensory context *s*_*i*_ ∼ *ρ*(·). The learner samples a set of policy parameters *θ*_*i*_ from its belief *q*_*i*_(*θ*) over policies, and it acts according to 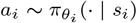. It then receives reward *r*_*i*_ = *R*(*s*_*i*_, *a*_*i*_) and updates its belief. **B)** With experience, the belief converges from the prior policy distribution *q*_0_ toward the optimal policy distribution *q*_*\**_ demanded by the task. The Kullback–Leibler (KL) divergence between the two distributions, *D*_KL_(*q*_*\**_ ∥ *q*_0_), measures the alignment between the prior and the task demand (larger values indicate poorer alignment). We call this the Effective Complexity (EC, Φ) of the task for this learner. **C)** Different tasks can demand different optimal policies, schematized here as local points in the space of all policies. For a learner whose prior is equidistant from both task-optimal policies, the shape of the prior determines the relative learning speed on the two tasks. Under a uniform prior (left), both tasks are learned at the same average speed. Under a nonuniform prior (right), the task aligned with the high-probability direction of the prior (task 1) is learned faster. **D)** Predicted learning curves for task 1 (green) and task 2 (blue) under the nonuniform prior. Our theory predicts a lawful relationship between the learning speeds of different tasks. In particular, the number of trials needed to reach the same level of performance Δ(dashed lines) is proportional to the effective complexity Φ. Therefore, effective complexity multiplicatively sets the speed of learning.

The goal of the agent is to use trial-to-trial feedback to gradually learn an optimal policy distribution *q*_*_ that maximizes performance. We measure this performance by regret, defined as the difference between the maximum reward and the received reward (the maximum reward is used only in our analysis and need not be known by the learner; see *Methods*).

We assume that the agent learns through an iterative Bayesian update rule,

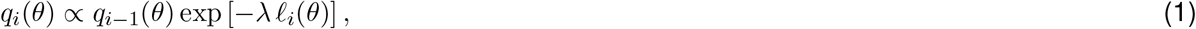

where *ℓ*_*i*_(*θ*) is an estimate of the regret that each policy *θ* would have incurred on trial *i* based on the outcome *r*_*i*_, and *λ >* 0 sets how strongly each trial updates the belief, analogous to a learning rate (see *Methods*). We assume that the agent begins with a prior *q*_0_(*θ*). Over the course of learning, the agent’s belief will approach the optimal policy *q*_*_(*θ*) (Fig. 1B) that achieves minimal regret on the task (which need not be zero). The distribution *q*_*_(*θ*) can be viewed as describing the task demand. Note that we do not need to specify the form of the policies or of these distributions; we only assume that they exist.

Conceptually, the prior distribution encodes the agent’s inductive bias; i.e., the assumptions that the agent brings to the task about which stimulus–action associations are more likely to lead to reward. Our main result is a quantitative description of how the learning speed depends on the alignment between this inductive bias and the task demand. Let *q*_*m*_ denote the learner’s policy distribution after *m* trials of learning. We derive an upper bound on Δ(*m*), the average gap in regret between *q*_*m*_ and *q*_*_. Specifically, given a Bayesian learner with prior *q*_0_ and task demand *q*_*_, the gap Δ(*m*) satisfies (see *Methods*):

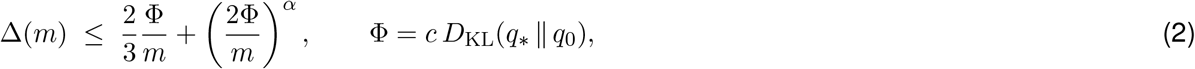

where 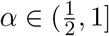 and *c* ≥ 1 are constants for a given learner and task.

We call Φthe Effective Complexity (EC) of the task for the learner. It is set by the Kullback–Leibler (KL) divergence between the prior *q*_0_ and the optimal policy *q*_*_, which measures how well the agent’s inductive bias aligns with the task demands (Fig. 1B). The KL divergence can be expressed in bits, and interpreted as the cost of representing the optimal policy *q*_*_ under the animal’s prior *q*_0_ (see *Methods*).

Larger values of EC indicate poorer alignment and result in slower learning. For example, if an agent maintains a uniform prior over all possible policies, then any two tasks will be learned at the same average speed (Fig. 1C, left). However, if an agent maintains a nonuniform prior whose probability density better aligns with the optimal policy of one task, then the aligned task will be quicker to learn, and the unaligned task will be slower to learn (Fig. 1C, right). Thus, it is not the apparent complexity of the task itself, but rather how well the task is aligned with the prior, that determines the speed of learning.

More importantly, our theory makes a precise prediction about learning speed. A typical account would attribute the difference between fast and slow learning to different learning rates, for example, by fitting a model to the learning curves [34, 35]. In our theory, by contrast, performance Δ(*m*) depends on experience only through the ratio Φ*/m*. Thus, effective complexity *multiplicatively* scales the number of trials required for learning. This means that a task with twice the effective complexity requires twice as many trials to reach the same performance (Fig. 1D). Unlike a fitted learning rate, this scaling factor is set by the alignment between the prior and the task, and can therefore be predicted *before learning begins*. Also note that the entire influence of the prior and task demands on the speed of learning is summarized by a single number. Therefore, two learners with very different priors should learn a task at the same speed if the task has the same effective complexity for both.

### Effective complexity predicts learning speed

The theory makes a strong prediction: if the prior can be empirically measured *before learning begins*, the effective complexity can be calculated for a given set of tasks, and should predict the relative learning speed between the tasks *without any free parameters*. Because the effective complexity (Φ) multiplicatively scales the number of trials (*m*) required to reach a given performance level (Eq. 2), log *m* should increase with log Φwith a slope of exactly one (Fig. 2A; see *Methods*).

**Figure 2:**
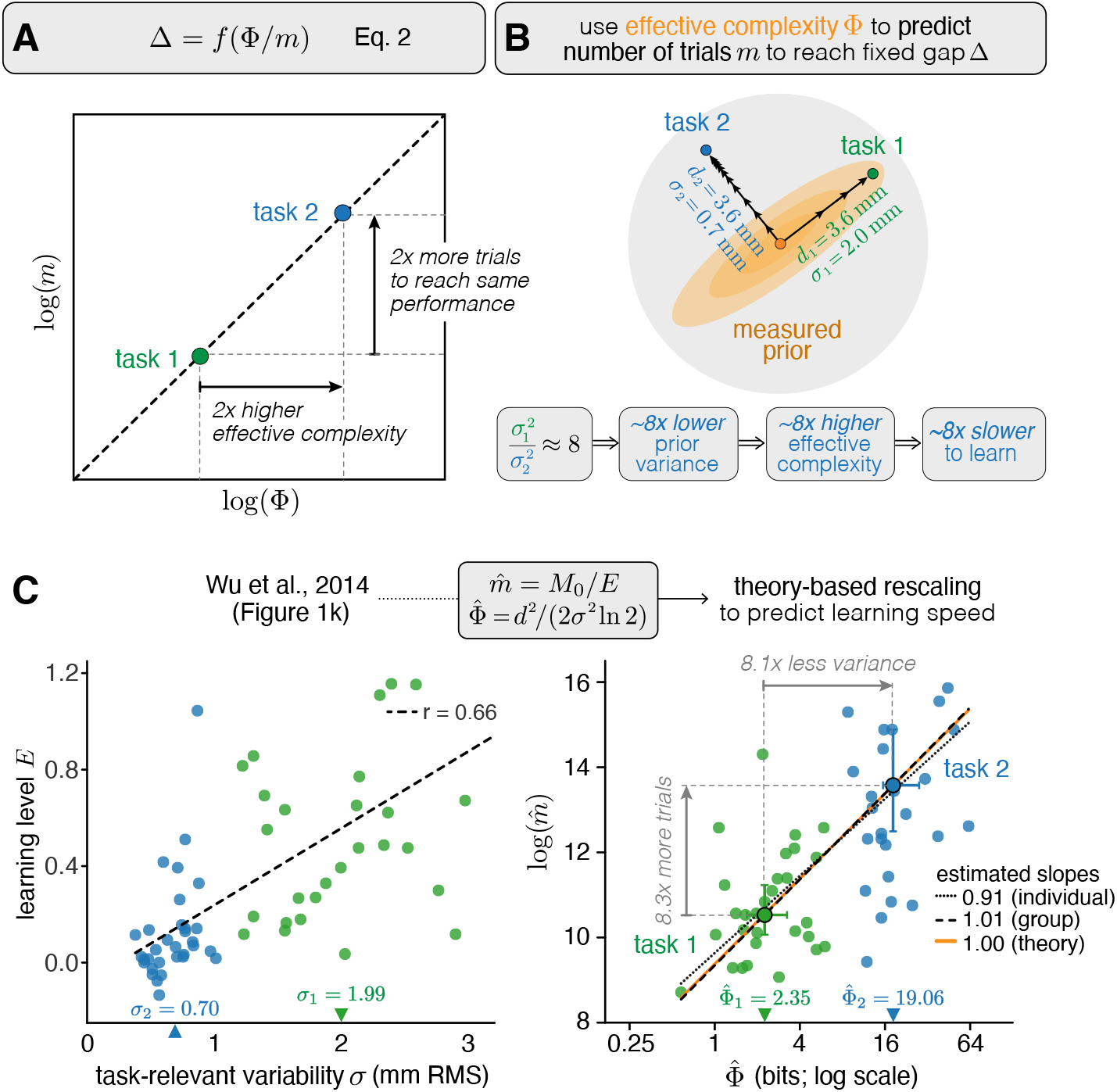
Effective complexity measured before learning predicts learning speed. **A)** Schematic. Because the performance gap Δis a function of the ratio Φ*/m* (Eq. 2), log *m* and log Φare linearly related with a slope of 1. As a result, if one task has twice the effective complexity than another, theory predicts it will require twice as many trials to reach the same performance. **B)** Schematic. We consider two tasks that are defined relative to a prior distribution that was estimated from baseline movement variability (orange). Both tasks require a change in movement of the same magnitude (*d* = 3.6 mm RMS), but task 1 (green) is aligned with a direction of high baseline variability (*σ* = 2.0 mm), whereas task 2 (blue) is orthogonal to it (*σ* = 0.7 mm). Under a Gaussian prior, effective complexity is inversely proportional to variance *σ*^2^. Therefore, task 2 (blue) has approximately 8 times higher effective complexity, and will thus be approximately 8 times slower to learn. **C)** Left: average learning level *E* of human participants over the *M*_0_ training trials as a function of task-relevant baseline variability *σ*, reproduced from Wu et al. [36], Fig. 1k. Each dot is one participant; dashed line: linear fit across all participants. Right: the same data after theory-based rescaling, showing the estimated number of trials to proficiency, 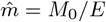, against the measured effective complexity, 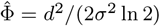, both on a log scale. Orange line: theoretical prediction (slope = 1). Black dashed line: fit to the task medians (large circles; error bars: bootstrapped 95% CI). Dotted line: linear fit across participants. The median 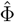 was 2.35 and 19.06 bits for task 1 and task 2, respectively (8.1 times higher for task 2); equivalently, the median variance along the task 2 direction was 8.1 times lower. The median estimated number of trials was 8.3 times higher for task 2.

To test this prediction, we reanalyzed data from a study by Wu et al. [36]. In this study, human participants used a handle to make rapid reaching movements without visual feedback; after each trial, they received only a numerical score reflecting how closely their movement matched a hidden target trajectory. Before learning, baseline movement variability was measured for each participant. This baseline variability provides an estimate of the prior distribution over movement policies that participants bring to the task. Participants were then assigned to one of two learning tasks: in task 1, the hidden target was aligned with a direction of relatively high baseline variability, while in task 2, the hidden target was orthogonal to it. In both tasks, the required change in movement had the same magnitude *d* (measured as root-mean-square, RMS, units in mm; Fig. 2B). The original study found that variability along each task direction (*σ*; RMS deviation from the average trajectory) was positively correlated with the average learning level *E* over the first *M*_0_ training trials (Fig. 2C, left).

Assuming a Gaussian prior over movements centered on the average movement trajectory, our theory allows us to estimate the effective complexity of the task, 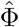, from the movement variability 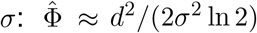 . In other words, for a fixed change in movement *d*, the effective complexity is proportional to 1*/σ*^2^. Our theory therefore explains why a target aligned with the direction of high movement variability was faster to learn, because the larger *σ* along that direction implies a lower effective complexity. We further estimated the number of trials needed to reach proficiency as 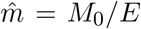, and found that 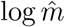 and 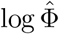 were linearly related with a slope close to the predicted value of 1, both across participants (slope = 0.91, 95% CI [0.59, 1.22]) and between task averages (slope = 1.01). This relationship held when 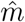 was estimated under two alternative assumptions (Fig. S1). The data also illustrate the multiplicative effect of EC: The median effective complexity was 2.35 bits for task 1 and 19.06 bits for task 2, an 8.1-fold increase that closely matched the 8.3-fold increase in the median estimated number of trials needed to learn (Fig. 2C, right). Thus, by estimating EC from baseline behavior before learning, it is possible to precisely predict the relative learning speed of the two tasks.

### Learning trajectories follow the predicted scaling law

The previous results show how this theory enables us to predict overall learning speed from baseline behavioral variability. The theory makes additional predictions about the temporal evolution of performance during learning. For example, by slightly relaxing Eq. 2, taking the logarithm of both sides, and absorbing the constant prefactor into a rescaled Φ^′^ (see *Methods*), we can rewrite the gap to optimal performance as:

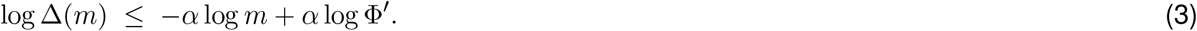

This reveals a linear relationship between log Δ(*m*) and log *m*. In other words, the gap to optimal performance decreases as a *power law* in the number of trials. The constant −*α* sets the slope of the learning curve, and Φ^′^ sets its intercept. Geometrically, this means that a lower effective complexity speeds learning by shifting the learning curve to the left in log–log coordinates (Fig. 3A); i.e., the multiplicative scaling of effective complexity becomes an additive shift in intercept.

**Figure 3:**
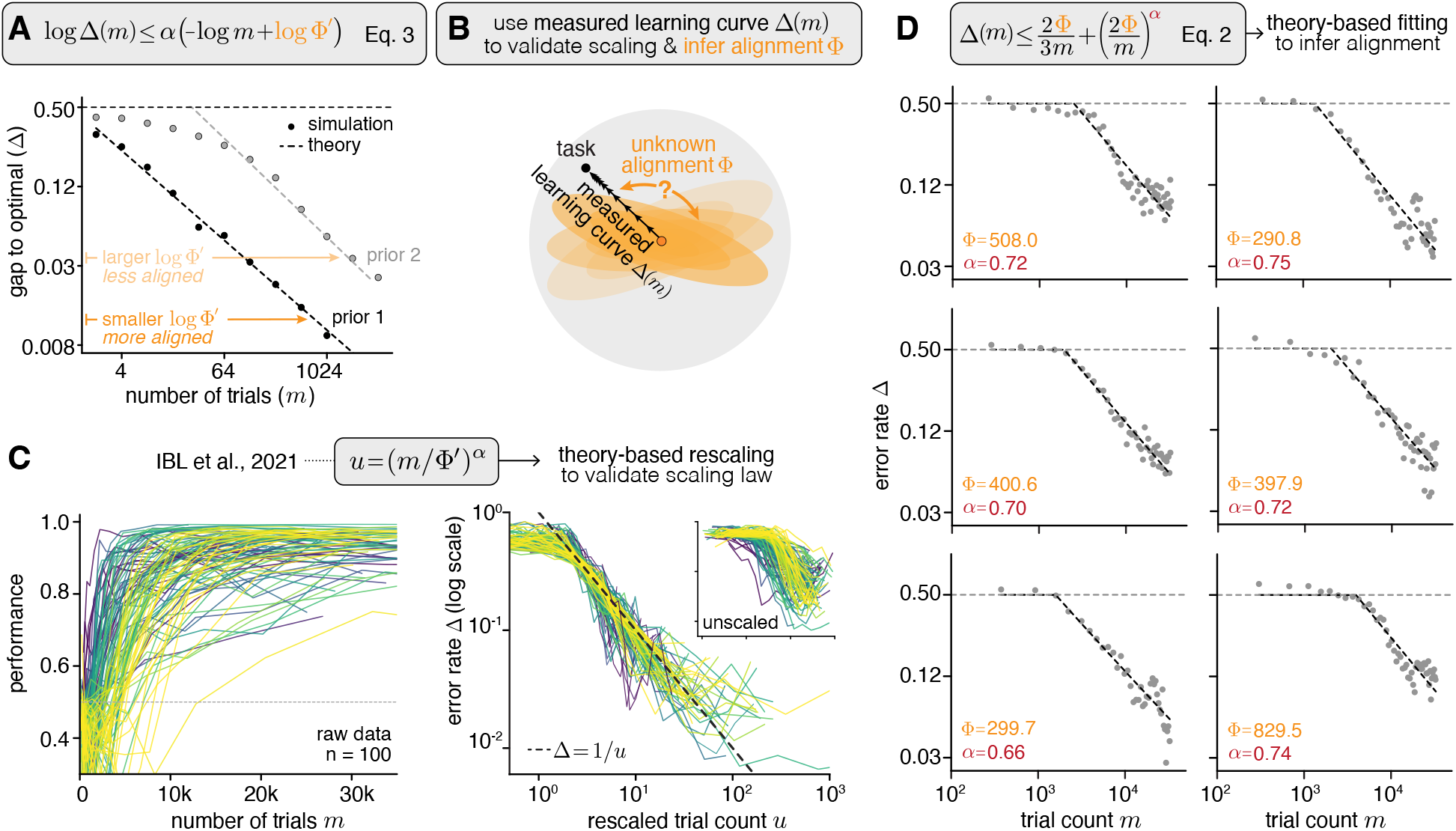
Individual learning curves collapse onto a single function predicted by the scaling law. **A)** Theory predicts that the average gap Δbetween the learner’s performance and optimal performance decreases as a power law in the number of trials *m*. In log-log coordinates, this is a linear function with its intercept determined by the effective complexity Φ. Given a fixed task, a learner whose prior is more aligned to the task (darker orange) will have a learning curve shifted to the left relative to a learner whose prior is less aligned (light orange). **B)** Schematic. When an animal’s prior is unknown, it is possible to use measured learning curves to validate the scaling law and to infer the alignment between the prior and the task. **C)** Individual learning curves from the IBL dataset [3] collapse onto a single function predicted by the theory after rescaling. Left: individual learning curves for *n* = 100 mice, displayed in their typical format. Right: the same learning curves, replotted as error rate versus trials in log–log coordinates, with trial count rescaled as *u* = (*m/*Φ^*′*^)^*α*^. Inset: error rate curves in log–log coordinates without rescaling. Individual mice were selected for display based on the smoothness of their learning curves; see Supplementary Fig. S2 for all mice. **D)** The average learning curves for each laboratory cohort (gray markers) are well fit by the two-term bound (black-dashed line; Eq. 2). The best-fitting values of Φcan be used to infer the effective complexity of the IBL task. Note that one of the seven laboratories is not shown, because many of its mice performed above chance from the beginning of the recorded sessions (see Supplementary Fig. S3A). Also see Supplementary Fig. S3C for the estimated parameters from individual mice.

The power-law improvement in performance predicted by Eq. 3 is similar to the data scaling laws observed in large machine learning models [14]. To test whether this scaling law also describes biological learning, we reanalyzed the International Brain Laboratory (IBL) dataset [3], in which cohorts of mice across seven laboratories were trained to perform a standardized visually-guided decision task. In this task, head-fixed mice were placed in front of a virtual screen that displayed visual gratings of high or low contrast. On each trial, a single grating appeared on either the left or right side of the screen, and mice learned to turn a wheel to move the grating to the center of the screen for a water reward. We measured learning by the error rate on high-contrast trials, for which the optimal error can be assumed to be close to zero.

Although in this setting, the animals’ priors are unknown, the scaling law makes a prediction that can be tested without them (Fig. 3B): if the bound is tight, rescaling the trial count for each animal individually as *u* = (*m/*Φ^′^)^*α*^ should collapse all learning curves onto a single function. Indeed, we found that applying this rescaling, with Φ^′^ and *α* estimated for each animal, collapsed the learning curves onto Δ= 1*/u*, as predicted (Fig. 3C).

Beyond testing its functional form, the scaling law allows us to decompose each learning curve into two interpretable components: the effective complexity Φ, and the exponent *α*. To infer these components, we fit the full two-term bound (Eq. 2) to the average learning curves of each laboratory (see *Methods*). These fits yielded consistent estimates of both Φand *α* (Fig. 3D), suggesting that the effective complexity is a reproducible property of the task for different cohorts of mice. Fits to individual mice corroborated this conclusion, with a similar median EC and median exponent to those observed across labs, and with the majority of exponent estimates falling close to the theoretical range (Supplementary Fig. S3C). Finally, the finding that the best-fitting EC values are large in magnitude suggests that the nominally simple IBL task is poorly aligned with the prior distributions of the mice, and therefore requires extensive experience to learn (see *Discussion*).

## Discussion

Here, we combined a theoretical analysis of a Bayesian learner with empirical analyses of data from humans and mice to show that biological learning follows a universal scaling law. This scaling law states that learning exhibits a power law dependence in *m/*Φ, the number of trials normalized by the effective complexity of the learning task. This is a statement about proportionality: within a given task, doubling the amount of experience reduces the gap to optimal performance by a constant factor; across tasks, doubling the effective complexity doubles the amount of experience required to reach the same level of performance. This scaling law offers an explanation for why learning speed does not reflect nominal task complexity, but instead depends on how well the demands of the task are aligned with the inductive biases of the animal. By formalizing inductive bias at the level of behavioral policies, we showed that it can be measured from baseline behavior and used to predict subsequent learning speed.

The idea that performance improves as a power law in experience was first proposed by Newell and Rosenbloom [30] as the “power law of practice”, used to describe human learning performance. However, its generality has been debated, with some accounts suggesting that it is an artifact of averaging over individual learning curves that are themselves exponential [31, 37]. Our theory offers a new perspective on this debate, by showing that this power-law dependence is evident in other animals, beyond humans, at the level of both average and individual learning curves. Demonstrating this dependence relies on two methodological points. First, whereas the classic result primarily concerns reaction times, our theory concerns the gap to optimal performance, a quantity that the learning is putatively aiming to minimize. Second, because the power law applies to the gap, the asymptotic performance must be known or estimated; otherwise, a learning curve that follows a power law in principle can appear to deviate from one in practice. By analyzing high-contrast trials in the IBL dataset, for which the optimal error is close to zero, we avoid this ambiguity. This allowed us to demonstrate that our two-term bound describes individual learning curves better than alternative functional forms, including exponential and sigmoid functions (Supplementary Fig. S3B).

In our theory, the effective complexity can be interpreted as the number of bits required to represent the optimal policy under the learner’s prior. This connects our work to the literature on minimum description length learning [38, 39]. For example, Feldman [21] showed that people are slower to learn logical rules with higher Boolean complexity. This connection highlights a subtle but important distinction between complexity and alignment. In the Boolean complexity measure, learning difficulty is a fixed property of the rule, because the description length is computed in a fixed logical language, which implicitly assumes a prior that assigns higher probability to simpler rules. In our formulation, this need not hold. A learner can assign high probability to a small subset of complex rules, so that learning those rules is faster, at the cost of slower learning for other rules. What determines learning speed is therefore not the complexity of the rule itself, but its *effective* complexity under the learner’s prior. We think this distinction is particularly important for thinking about animal learning, because we tend to judge task difficulty by our own prior over what is simple, which might not match the prior of the animal.

Our theory also allows the effective complexity of a task to be inferred from the shape of the learning curves, as we demonstrated for the IBL task. Assuming a tight bound with the constant *c* of order one, the median KL divergence of the IBL task is roughly 500 bits. This is an order-of-magnitude estimate (see *Methods*); nevertheless, it is far greater than the 2 bits needed if mice were learning a simple mapping between two stimuli (a left versus right grating) and two actions (a left versus right turn) under a uniform prior [1]. This comparison offers a concrete interpretation of the misalignment between the prior and the task, and suggests that learning does not take place in the four-hypothesis space of matching turn directions to grating locations. Instead, mice must discover which aspects of the stimulus, their motor actions, and the associations between them are relevant to the task, which together form a much larger space in which learning occurs.

Our work formulates inductive bias at the level of behavioral policies, which is complementary to formalisms based on neural representations, such as kernel alignment [17] and population geometry [40]. The two are nonetheless closely related. For example, assuming a Gaussian prior over motor policies in the human motor learning task, a target that shifts the movement by *d* along a direction *u* has an effective complexity that is proportional to *d*^2^ *u*^⊤^Σ^−1^*u*. This quantity weights the target by the inverse of the prior covariance, such that movements along high-variance directions have low effective complexity and are learned from fewer trials. This is closely analogous to spectral bias in kernel regression, where target functions aligned with high-eigenvalue directions of the kernel are learned faster [17, 19]. When combined, these frameworks could offer a common language for relating inductive bias across behavior, neural representations [41], and circuit architectures [42, 43].

Several caveats apply to our results. First, our analysis focused on the simplest form of the bound on the performance gap Δ, using the optimal policy distribution. This form is not the tightest, and the predicted power law is expected to hold mainly at large trial counts; as a simple example, performance at small *m* is bounded by chance. Our theoretical analysis does yield a tighter bound that may better describe the initial phase of learning (see *Methods*), and we leave its investigation to future work. Second, the KL divergence between the prior and the task demand determines the effective complexity only up to a scale factor that depends on the reward structure and on how the learner estimates reward. However, this factor should be approximately constant across tasks that share the same format, and therefore cancels when comparing them. In the human motor learning task, for example, the two tasks differed only in the direction of the target, and the ratio of their estimated KL divergences predicted the ratio of their learning speeds, as expected from the theory. How well the same quantity predicts learning speed across different contexts, and at what level of description the prior and the task demand should be measured, remain open empirical questions.

### Outlook

Much of the recent progress in artificial intelligence has been driven by scaling laws: building systems that are large enough and flexible enough to achieve low generalization error once sufficient data and compute are invested [14]. Our results suggest that biological learning also follows a data scaling law, but we speculate that it operates in a different regime. Rather than relying on weak inductive biases and abundant data, animals bring strong inductive biases, shaped by evolution and tailored to their ecological niche, that enable highly efficient learning in natural settings [44]. When placed in artificial settings, however, these same inductive biases appear as inefficiencies. What distinguishes biological learning, then, may lie not in the exponent of the scaling law, but in its intercept.

## Acknowledgements

This work was supported by the Howard Hughes Medical Institute.

## Methods

### Theoretical Framework

#### Multi-armed Contextual Bandit

We begin by framing a typical trial-based laboratory learning task as a contextual bandit problem [33], and we model an animal in the task as a Bayesian learning agent. On each trial *i*, the environment generates a sensory context *s*_*i*_ ∼ *ρ*(*s*), where *ρ*(*s*) is unknown to the agent. The agent selects an action *a*_*i*_ ∈ *A* and receives feedback *r*_*i*_ = *R*(*s*_*i*_, *a*_*i*_) according to a fixed reward rule.

We assume that the agent’s behavior is described by a set of policies *π*_*θ*_(*a* | *s*), parameterized by *θ* ∈ Θ, where Θ is a measurable parameter space. Rather than committing to a single policy, the agent maintains a policy distribution *q*_*i*_(*θ*) over Θ. On each trial, the agent samples a policy parameter *θ*_*i*_ ∼ *q*_*i*_(·), and then samples an action according to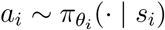.

After *m* trials, we denote the observed experience by *S*^*m*^ = {(*s*_1_, *a*_1_, *r*_1_), …, (*s*_*m*_, *a*_*m*_, *r*_*m*_)}. We use *D*^*m*^ to denote the distribution over the length-*m* sequences induced by the contextual bandit and the agent’s learning dynamics (i.e., *S*^*m*^ ∼ *D*^*m*^).

We note that in general, the space of sensory inputs *S* and actions *A* need not (and most likely will not) correspond directly to the experimenter-defined task structure; rather, they should be understood as the spaces over which learning occurs from the animal’s perspective.

We assume there is a maximum attainable reward 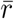, and we define the per-trial regret Δ_*i*_ as 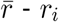. The expected (true) regret [45] of a policy *π*_*θ*_ is:

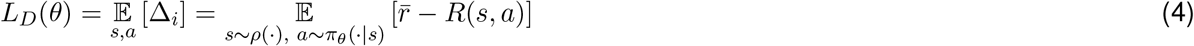

Note that we use the regret formulation for consistency with the bandit literature, but this is only a matter of convention: since regret and reward differ by a constant 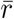, minimizing expected regret is equivalent to maximizing expected reward, and the constant cancels out in the derivation. 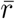 need not be known or specified.

In addition, for any distribution *q*(*θ*), with slight abuse of notation, we define the expected regret of a behavioral policy distribution as:

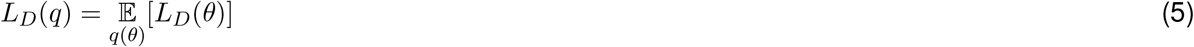

The goal of the agent is to learn a policy distribution that minimizes the expected regret *L*_*D*_(*q*) (i.e., maximize the expected reward). In the next sections, we model the animal as a Bayesian learning agent and analyze its behavior.

### A Bayesian Learner

We now specify the learning rule by which the agent updates its distribution over behavioral policies. Prior to learning, we assume that the agent has a prior distribution *q*_0_(*θ*) over Θ. This prior acts as an *inductive bias*, capturing the agent’s prior knowledge and assumptions *before* any task-specific experience.

We assume that learning occurs through the construction of some empirical estimates of the expected regret based on the agent’s experience *S*^*m*^, defined as *L*_*S*_(*θ* | *S*^*m*^). For example, *L*_*S*_(*θ* | *S*^*m*^) can be the inverse propensity score estimator (see *Supplementary Proofs* for additional discussion on other valid estimators):

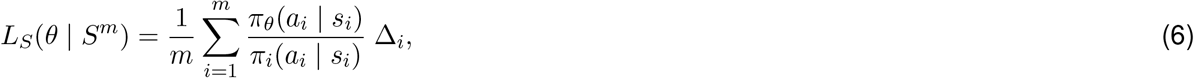

where *π*_*i*_ is the policy from which the agent samples an action on trial *i*.

After observing *m* trials, the agent computes a posterior distribution over policies according to a generalized Bayesian inference rule (the Gibbs posterior) [46, 47]:

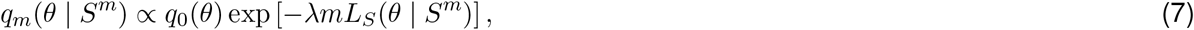

or according to its online equivalent:

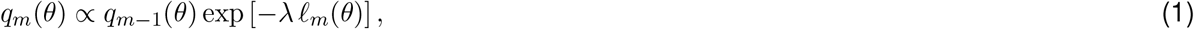

where *λ* is an inverse-temperature parameter, and *ℓ*_*m*_ is the single-sample equivalent of *L*_*S*_, computed using the sample at trial *m*.

We interpret *q*_*m*_(*θ* | *S*^*m*^) as the agent’s updated behavioral policy distribution after *m* trials. On trial *m*, the agent samples a parameter *θ* ∼ *q*_*m*_(·) and acts according to the corresponding policy *π*_*θ*_(*a* | *s*). For notational simplicity, we will write *q*_*m*_(*θ*) in place of *q*_*m*_(*θ* | *S*^*m*^).

For an agent that follows the learning rule defined above, its average learning curve as a function of trials is described by:

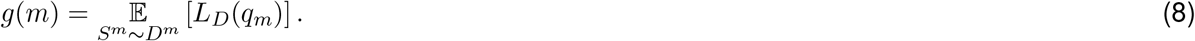

Lastly, let *q*_*_(*θ*) denote the “optimal” posterior that describe the agent’s behavior at the end of learning; *q*_*_ can be thought of as describing the task demands. In tasks where the agent becomes proficient, we would expect *L*_*D*_(*q*_*_) ≈ 0 (but it need not be zero in general). We denote the gap between *g*(*m*) and *L*_*D*_(*q*_*_) as

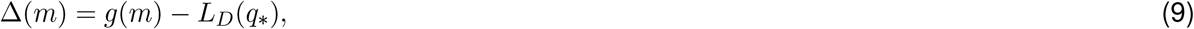

which is a function of *m*.

### Scaling Law for Biological Learning

Our main theoretical result is an upper bound on the average learning curve (see *Supplementary Proofs* for the full derivation). In particular:

*Given a contextual bandit setup with a reward function R*(*s, a*) *and a stimulus distribution ρ*(*s*), *consider a Bayesian learner with prior q*_0_(*θ*) *as described above. Then there exist fixed constants* 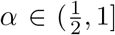 *and c* ≥ 1, *such that the average learning curve g*(*m*) *satisfies:*

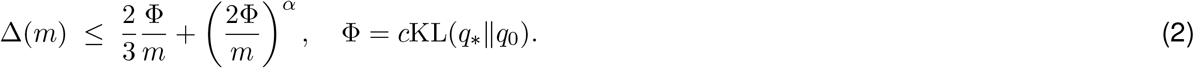

Here *α* is a rate parameter determined by how the variance of the regret estimate changes during learning, and *c* is a fixed constant for a given learner and reward scale. This equation delineates two factors that shape performance during learning: the amount of experience, captured by the number of trials *m*, and the agent’s inductive bias, captured by the prior-dependent KL term.

This equation also characterizes the rate of learning. Specifically, the expected regret of the learned behavior *L*_*D*_(*q*_*m*_) converges to that of the optimal behavior *L*_*D*_(*q*_*_) at rate *O*(*m*^−*α*^). Since *α* ∈ (1*/*2, 1], the bound predicts a power-law scaling [48, 49] with an exponent between −0.5 and −1.

To highlight the predicted power-law behavior, we can rewrite Eq. 2 using a slightly relaxed bound:

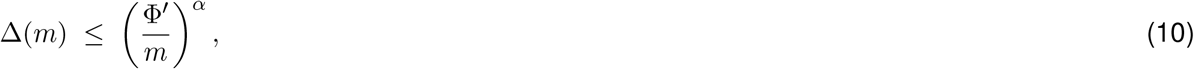

by absorbing the constant into 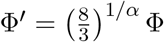. Taking the logarithm of both sides gives

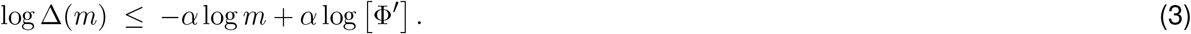

This reveals a linear relationship between log Δ(*m*) and log *m* in log–log coordinates, where −*α* sets the slope of the learning curve, and Φ= *c*KL(*q*_*_∥*q*_0_) sets its intercept (Fig. 3).

Lastly, the gap-to-optimal bound we presented in the main text (Eq. 2) is the easiest to interpret, but it is not the tightest, and it is expected to describe learning mainly at large *m*. For example, at small *m*, the bound exceeds chance-level error; in the data analysis, we therefore model the learning curve as the minimum of chance-level error and the bound. Our theoretical analysis, however, yields a tighter bound that may better describe the initial phase of learning (see *Supplementary Proofs*, Infimum Oracle Bound); we leave its detailed investigation to future work.

### Effective Complexity

We refer to the term Φ= *c*KL(*q*_*_∥*q*_0_) as the *Effective Complexity* (EC) of the task. This terminology [50, 51, 52] is meant to connect to classical statistical learning theory, where a fixed capacity measure, such as the VC dimension [53, 54], controls the sample complexity of the learning problem. The effective complexity, on the other hand, depends on both the agent’s prior *q*_0_(*θ*) and the task’s optimal solution *q*_*_(*θ*), and captures the *alignment* between them. When *q*_*_ is well aligned with *q*_0_, the task has low effective complexity and is easier to learn; conversely, learning a *q*_*_ that is poorly aligned with *q*_0_ requires more experience. Thus, it is not the task’s apparent complexity *per se*, but rather how well the target behavior is supported by the prior, that determines the speed of learning.

More precisely, notice that Δ(*m*) depends on experience through only the ratio Φ*/m*. Therefore, EC sets the number of trials required to reach a given level of performance (i.e., learning speed) *multiplicatively* : Consider two tasks with optimal policies 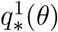 and 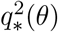, with 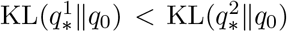. Let *m*_1_ and *m*_2_ denote the number of trials required for the animal to reach the same expected performance. Assuming the constants *α* and *c* are the same in both tasks, we obtain:

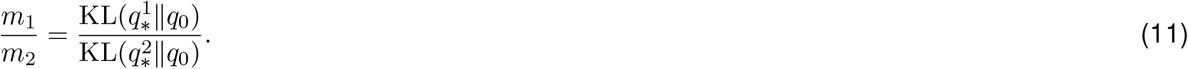

Therefore, the theory predicts that learning the more aligned policy 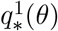 requires fewer trials by a precise fraction of 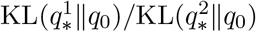, that is, the ratio of the effective complexities.

Equivalently, taking logs of both sides of Eq. 11, we have, for all pairs of *m* and *q*:

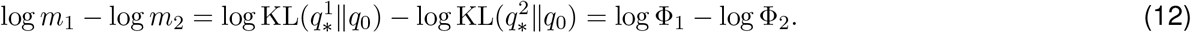

Hence, log *m* − log Φ= *k*; i.e., log *m* is linearly related to log Φwith a slope of exactly one (Fig. 2A).

Lastly, effective complexity is measured in nats (1 nat = 1*/* ln 2 ≈ 1.44 bits). This unit follows from the derivation naturally, in which the KL divergence arises from the logarithm of an exponential moment (Eq. 25). The KL divergence can be interpreted as the cost of representing the optimal policy distribution *q*^*^ under the learner’s prior [38, 39]. Eq. 11 can thus be read more intuitively: to reach a given level of performance, the number of trials needed is proportional to the number of bits of the task, measured by effective complexity.

## Data Analysis

### Human Motor Learning

We reanalyzed Experiment 2 of Wu et al. [36]. In this experiment, participants made rapid reaching movements without visual feedback of their hand path. A baseline block measured each participant’s average hand path and its trial-to-trial variability. In the training block, the only feedback participants were given was a numerical score for deflecting their hand path toward a hidden target deflection, which had the same magnitude in both tasks (*d* = 3.6 mm RMS) but different shapes: along a direction of high baseline variability in task 1 (*n* = 29), and along a direction of low baseline variability in task 2 (*n* = 32). The baseline variability of the hand path along each target direction is called the task-relevant variability *σ*, expressed in mm RMS. The average *σ* across participants is 2.0 mm for task 1, and 0.7 mm for task 2 (Fig. 2B). Learning on each trial was measured by projecting the hand path, relative to the baseline average, onto the target deflection. The learning level *E* is the average of this single-trial measure over *M*_0_ = 800 training trials.

### Effective complexity from baseline variability

We represent a movement policy by its hand path *θ* ∈ R^*k*^. Because the target is defined relative to the average baseline path, we take this average as the origin, and we model the prior as a multivariate Gaussian distribution over hand paths, *q*_0_(*θ*) = *N* (*θ*; 0, Σ), whose covariance Σ describes the baseline variability. The target of a task is described by its unit direction *w*, and the magnitude *d*.

In the simplest model of learning, the policy distribution keeps its shape and is shifted to a new location, *q*_*_(*θ*) = *N* (*θ*; *µ*, Σ). The KL divergence between two Gaussian distributions with the same covariance is

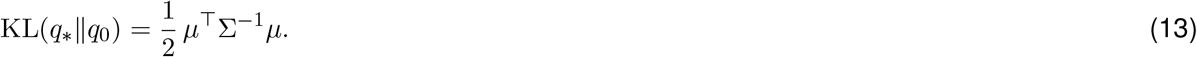

Any shift that satisfies *w*^⊤^*µ* = *d* solves the task equally well. Among these, we take the shift with the smallest KL divergence, which gives the tightest bound:

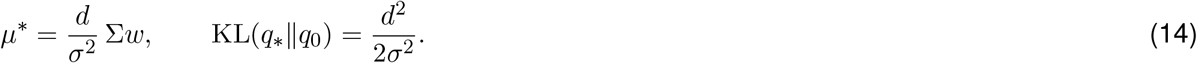

Therefore, the KL term is inversely proportional to the measured *variance, σ*^2^, along the direction of *w*. Intuitively, when *w* is an eigenvector of Σ, *µ*^*^ = *d***w** is a shift directly toward the target, where the learning cost of moving along that direction is inversely proportional to the variance *σ*^2^ along *w*.

Note that if we instead assume that the learned policy converges to a point mass at *µ*^*^, approximating the KL divergence by discretizing the space *θ* gives the same term *d*^2^*/*(2*σ*^2^), plus terms that do not depend on the target.

We therefore defined the measured effective complexity in this task, expressed in bits, as

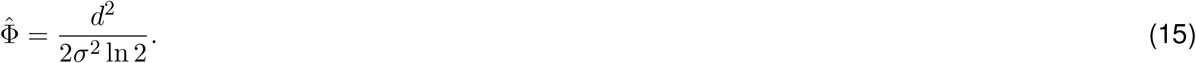

### Trials to proficiency

To compare the learning speed of human subjects with the predictions of the theory, we converted the learning level *E* into an estimate of the number of trials needed to reach proficiency (i.e., a fixed performance level),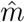.

In the simplest model, used in the main text, the learning level is inversely proportional to the number of trials needed to learn, so that 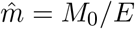. This model requires no fitting, and 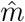 is a direct transformation of the measured *E*.

We also considered two alternative models of learning curves that saturate at an asymptote *A*: an exponential learning curve, 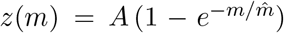, and a saturating power-law learning curve, 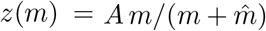, in which 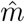 sets the number of trials needed to reach a fixed fraction of the asymptote. For each participant, we found the 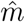 for which the average of *z*(*m*) over the first *M*_0_ trials equals the measured *E*. We set a common asymptote *A* = 2, just above the highest measured learning level (*E* = 1.72). The estimated slopes (see below) depended only weakly on this choice: for *A* between 1.8 and 100, they ranged from 0.91 to 1.19 for both models. All three models require *E >* 0. Five participants, all in task 2, had *E* ≤ 0, for which 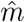 is not defined. They were excluded from the analysis of individual participants (leaving *n* = 56), but not from the group medians (see below).

### Testing the predicted scaling

The theory predicts that the number of trials needed to reach a given level of performance is proportional to the effective complexity, so that 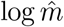 increases with 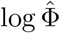 with a slope of one (Eq. 11; see *Effective Complexity*). Because the theory predicts the ratio of trials needed for the two tasks, the choice of units and the constant *c* do not affect this prediction.

In the motor task, this prediction takes a particularly simple form. Because the targets of the two tasks have the same amplitude *d*, 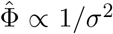, and the theory predicts that the ratio of trials needed to learn the two tasks equals the inverse ratio of their task-relevant baseline variances:

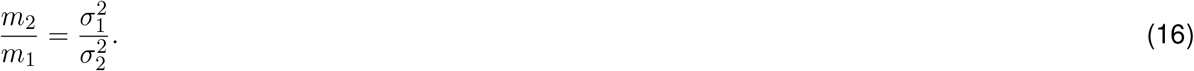

In other words, a target along a direction with *k*−times less baseline variance should require *k*−times more trials to learn (Fig. 2B).

We tested this prediction in two ways. Across participants, we regressed 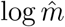 on 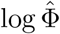 and estimated the slope. Second, between tasks, we computed the median *σ*^2^ and the median *E* of each task, converted them to 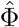 and 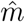, and computed the slope of the line through the two task medians across all participants. Confidence intervals for the task medians (95%) were obtained by bootstrap resampling within each task using 5,000 samples. The same analyses were repeated for all three models of the learning curve (Supplementary Fig. S1).

### IBL Learning Curve

We analyzed the training data of *n* = 140 mice from the International Brain Laboratory et al. [3] dataset. For each mouse and each training day *k*, we computed the error rate on high-contrast trials (contrast ≥ 50%), *y*_*k*_, and the cumulative number of trials completed by the end of that day, *m*_*k*_.

### Rescaling the learning curves

To rescale the learning curves, we fit each mouse’s learning curve with the power-law form of the bound (Eq. 10). We assumed that the optimal error on high-contrast trials is zero (*L*_*D*_(*q*_*_) = 0), so that Δ(*m*) equals the error rate, and we capped the predicted error at chance:

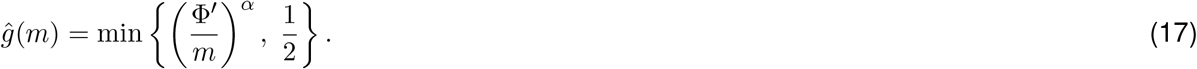

We estimated Φ^′^ *>* 0 and *α >* 0 by minimizing the squared error between the predicted and observed error rates, 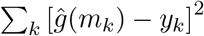, using bounded nonlinear least squares.

We fit the error rate on a linear, rather than logarithmic, scale in order to retain days with zero errors. The optimization was initialized from a straight-line fit in log–log coordinates, whose slope and intercept give initial values of −*α* and *α* log Φ^′^, respectively. Four mice whose fits explained less variance than a constant (*R*^2^ *<* 0) were excluded, leaving *n* = 136 mice.

Given the fitted parameters, we rescaled each mouse’s trial count as *u* = (*m/*Φ^′^)^*α*^. If the power law describes the individual learning curves, the rescaled curves should collapse onto the line Δ= 1*/u* in log–log coordinates (Fig. 3C).

Plotting daily error rates directly in log–log coordinates produces very noisy curves late in training, because each daily error rate is estimated from a limited number of high-contrast trials. For display purposes only (all fits were performed on the unbinned daily data), we therefore grouped the training days of each mouse into 15 bins evenly spaced in log *m* between its first and last day, and within each bin computed the trial-weighted average of the cumulative trial count and the error rate.

In addition, overlaying the noisy learning curves of all mice obscures their shape, so in Fig. 3C, we show the *n* = 100 mice with the smoothest learning curves, quantified as the average deviation of each log-error-rate from the mean of its two neighbors. This score is small for smooth curves, regardless of their shape relative to Δ= 1*/u*. The rescaled learning curves of all *n* = 136 mice, divided into three groups by smoothness, are shown in Supplementary Fig. S2.

### Estimating effective complexity

To more accurately estimate the effective complexity Φand the exponent *α*, we fit the full two-term bound (Eq. 2), assuming zero optimal error and capping the predicted error at chance:

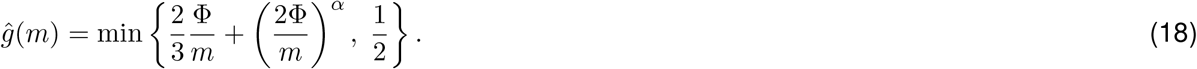

Φ*>* 0 and *α >* 0 were estimated by bounded nonlinear least squares on the error rate. The optimization was initialized from the same straight-line fit in log–log coordinates. We did not constrain *α* to the theoretical range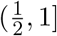; as a result, the fitted exponents provide a test of this prediction.

We fit Eq. 18 at two levels. First, for each laboratory cohort, we averaged the learning curves across mice by training day. Note that because mice were trained for different numbers of days, later days are averaged over progressively fewer mice; we therefore restricted the cohort fits up to *m* ≤ 2^15^ = 32,768 trials (Fig. 3D). The estimated parameters were consistent across six of the seven cohorts; the outlier cohort is shown in Supplementary Fig. S3A.

Second, we fit Eq. 18 to the daily data of each mouse individually, without truncation (Supplementary Fig. S3C). Fit quality was measured by *R*^2^, the fraction of variance in the daily error rate explained by the fit. We also tested two alternative two-parameter learning-curve models, the exponential function *ĝ*(*m*) = *a e*^−*m/τ*^ and the sigmoid function 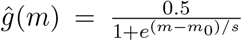. As with Eq. 18, both were capped at chance level during fitting. Our two-term function gave a higher *R*^2^ than both alternatives in 108 of 136 mice (Supplementary Fig. S3B).

As with the motor learning task, we convert the fitted Φinto bits. However, in this case, since we are not estimating KL, we need to know *c*. For a two-outcome task with bounded deterministic rewards, such as the IBL task, *c* ≤ 2 for an ideal learner, and larger for a noisier reward estimator (see *Supplementary Proofs*, Reward Estimator). The fitted Φtherefore provides an order-of-magnitude estimate of the KL divergence; for example, the median Φacross individual mice is approximately 500 bits (Supplementary Fig. S3C). Note that the estimation noise for Φis also logarithmic, since we are essentially relying on the intercept of the learning curves in log coordinates.

## Code Availability

All code used for the theoretical analysis, simulations, and data analyses in this paper is available at: https://github.com/lingqiz/inductive-bias.

## Supplementary Materials

### Supplementary Proofs

#### Proof of Eq. 2 and Eq. 10

In this section, we show how to arrive at the generalization bound [55, 46, 28, 29], as shown in Eq. 2 and Eq. 10. We begin by introducing a few additional definitions.

Define *ℓ*_*i*_(*θ*) as *L*_*S*_(*θ* | *S*^*m*^) (Eq. 6) estimated from a single sample (*s*_*i*_, *a*_*i*_, *r*_*i*_):

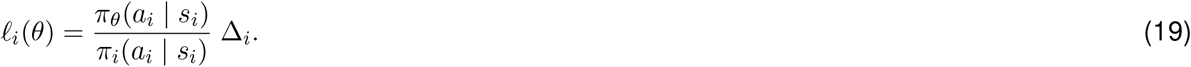

Similar to Eq. 5, we define *L*_*S*_(*q* | *S*^*m*^) = E_*q*(*θ*)_ [*L*_*S*_(*θ* | *S*^*m*^)], and *ℓ*_*i*_(*q*) = E_*q*(*θ*)_ [*ℓ*_*i*_(*θ*)]. As above, we use *q*_*_(*θ*) to refer to the optimal behavior at the end of learning. For a length-*m* sequence *S*^*m*^ ∼ *D*^*m*^, define the sequence of random variables:

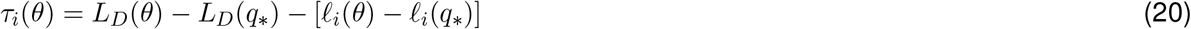

Note that the *τ*_*i*_’s are not i.i.d., because the actions *a*_*i*_ are generated by the agent based on the policy distribution *q*_*i*_ that is continuously updated through learning. We assume a bounded regret Δ_*i*_ ∈ [0, 1] and *τ*_*i*_ ≤ *b*. This can be achieved, for example, by normalizing the reward values and setting an exploration floor *η*, such that ∀*a* ∈ *A, π*_*i*_(*a*) ≥ *η >* 0. Lastly, we have E[*τ*_*i*_ | *τ*_1_, …, *τ*_*i*−1_] = 0, since we assume *L*_*S*_(*θ* | *S*^*m*^) is conditionally unbiased.

In this case, *τ*_*i*_’s form a martingale difference sequence. The cumulative 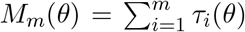 is a martingale [56], and its cumulative variance is defined as 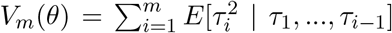. We then have:

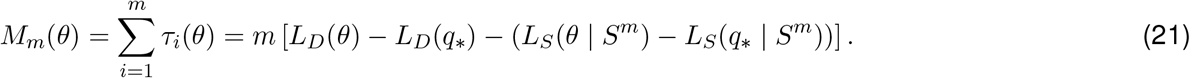

The quantity *L*_*D*_(*θ*) − *L*_*D*_(*q*_*_) is called the excess regret of policy *π*_*θ*_ (compared to *q*_*_). The first step of the proof is to show that for a fixed policy *θ*, the empirical estimate of the excess regret *L*_*S*_(*θ* | *S*^*m*^) − *L*_*S*_(*q*_*_ | *S*^*m*^) converges to the true excess regret at some rate.

We start with Freedman’s concentration inequality for martingale [57, 58, 59], which states that for any *t >* 0 and *v >* 0,

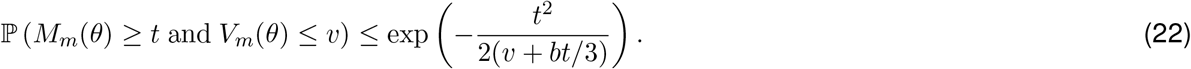

Equivalently, this concentration inequality follows from the exponential supermartingale bound:

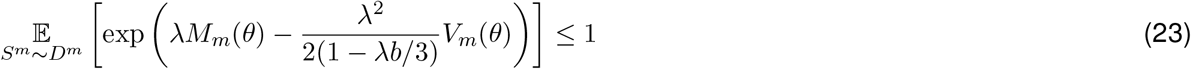

for any fixed *θ* and 0 *< λ <* 3*/b*.

The inequality holds for any fixed *θ* chosen independent of observing *S*^*m*^; in a learning problem, however, the learner uses *S*^*m*^ to choose (a distribution over) policies. Thus we need to lift the bound to a uniform statement that remains valid for any data-dependent choices of *θ*.

To this end, we define

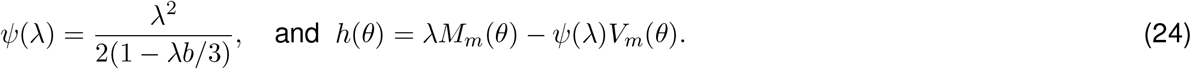

Taking the expectation on both sides of Eq. 23 with respect to the (data independent) prior distribution *q*_0_(*θ*), and exchanging the integral, we have:

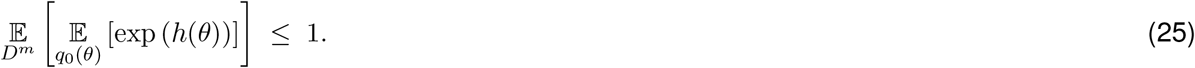

Taking the logarithm of both sides of the equation above, and expanding:

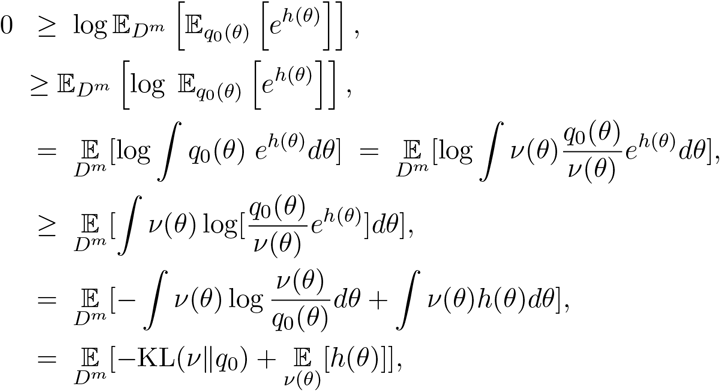

where both exchanges between logarithm and expectation use Jensen’s inequality. Inside of the expectation, this change-of-measure inequality holds for *any* distribution *ν*(*θ*) with finite KL(*ν*∥*q*_0_), including the ones that are *S*^*m*^ dependent. For each sample *S*^*m*^ ∼ *D*^*m*^, we may choose the posterior *ν*(*θ*) = *q*_*m*_(*θ* | *S*^*m*^) in evaluating the expectation while paying an extra cost of KL(*q*_*m*_∥*q*_0_).

We expand *h*(*θ*) and obtain:

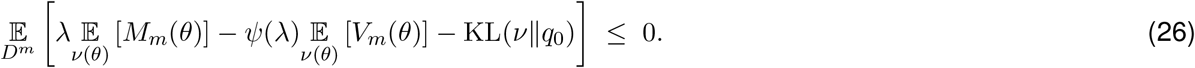

We abuse notation again and write 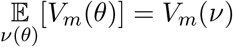. Plugging in Eq. 21 above, we have:

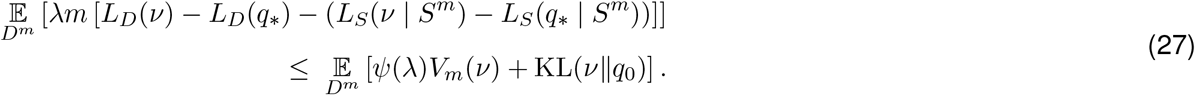

Rearranging the terms and dividing both sides by *λm* then gives:

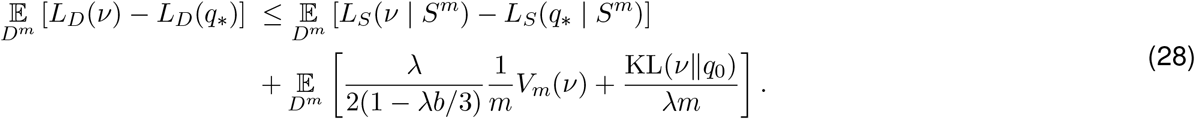

This is an upper bound on the true excess regret *L*_*D*_(*ν*) − *L*_*D*_(*q*_*_), expressed in terms of its empirical estimate *L*_*S*_(*ν*| *S*^*m*^) − *L*_*S*_(*q*_*_ | *S*^*m*^) and the complexity penalty KL(*ν*∥*q*_0_) for any *ν*(*θ*). Our next step is to show that, by choosing *ν*(*θ*) to balance empirical regret against the KL term, we obtain a learning rule with a guarantee on its true excess regret.

In particular, we notice that setting *ν*(*θ*) to the Gibbs posterior (Eq. 7):

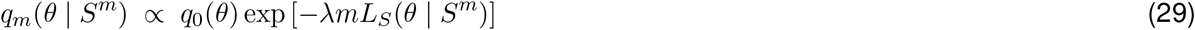

minimizes the quantity 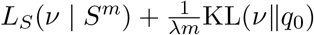 for any *S*^*m*^. Therefore,

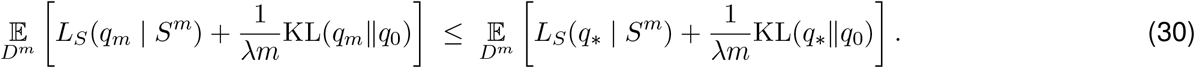

Substituting *ν*= *q*_*m*_ into Eq. 28, and plugging Eq. 30 into its RHS, we obtain:

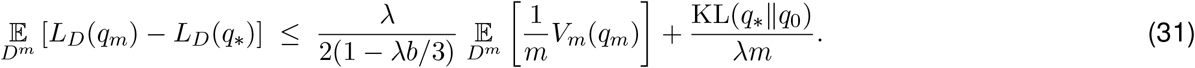

We have obtained an oracle bound of the average (over samples of *S*^*m*^) true excess regret of *q*_*m*_ compared to *q*_*_ that depends on *λ* and the variance term *V*_*m*_(*q*_*m*_).

Lastly, we introduce the Bernstein assumption [46, 60]:

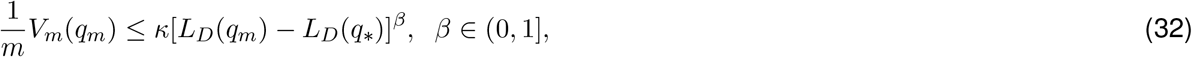

which relates the variance of the excess regret estimate for *q*_*m*_ to its value. The exponent *β* controls whether and how fast the variance shrinks as the excess regret becomes smaller: for *β* → 0, the variance only stays bounded by *κ*, whereas at *β* = 1, the variance decreases linearly with excess regret, such that the estimation becomes progressively easier as *q*_*m*_ approaches *q*_*_.

We therefore have:

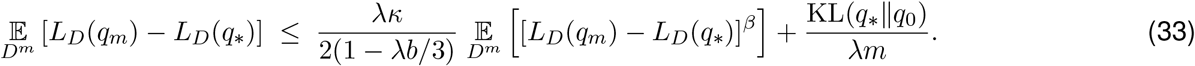

Let 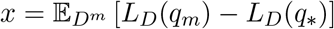. Since *L*_*D*_(*q*_*m*_) − *L*_*D*_(*q*_*_) ≥ 0, we apply Jensen’s inequality to the expectation on the RHS, and we move the exponent outside to arrive at the following inequality:

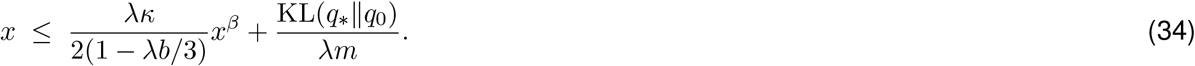

The last step of the proof is to solve this self-bounding inequality, which involves solving for the optimal sequence of *λ* that minimizes the RHS of Eq. 34. Define:

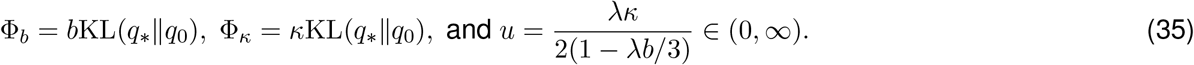

We then have:

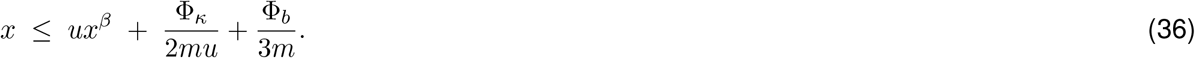

Let *A* = Φ_*κ*_*/*2*m* and *D* = Φ_*b*_*/*3*m*. We have (by assuming, on the RHS, that either the first term is larger or the remaining two terms are larger):

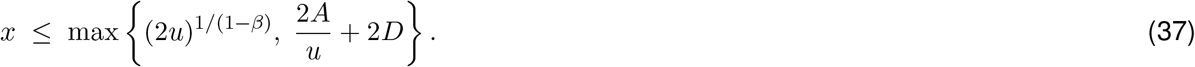

The first term increases and the second decreases as a function of *u*. We therefore choose the value of *u* (and hence *λ*) to balance the two sides by solving (2*u*)^1*/*(1−*β*)^ = 2*A/u*, which yields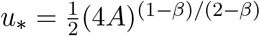. Substituting *u*_*_ back, we get:

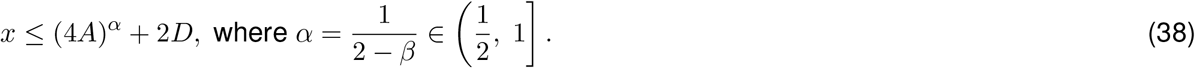

Note that the derivation above technically only holds for *β <* 1. However, for *β* = 1, we can simply set *u* = 1*/*2, and we recover the inequality above with *α* = 1.

The solution *u*_*_ specifies a sequence of inverse temperature values *λ*_*m*_ to attain the bound:

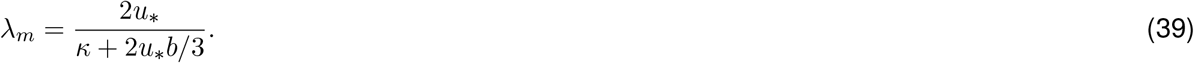

Lastly, by substituting the terms *A* and *D* into Eq. 38, we arrive at:

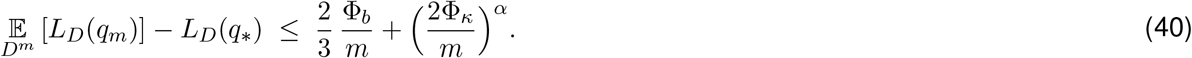

We simplify notation slightly by defining *c* = max(*b, κ*), and obtain our final result (Eq. 2):

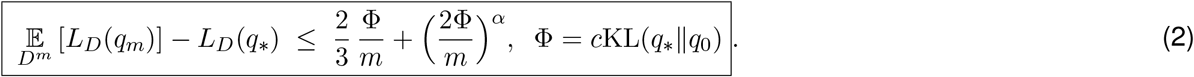

The rate parameter *α* controls the rate of convergence. When we can only assume bounded variance for the estimator, corresponding to *β* → 0, we obtain the slow rate *α* → 1*/*2. In contrast, when the variance shrinks linearly with excess regret, corresponding to *β* = 1, we obtain the fastest rate *α* = 1.

We can make additional simplifications to make the bound easier to interpret: for any *m* ≥ Φ, we have 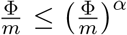, and 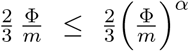. Combining the two terms on the RHS and absorbing the constant into 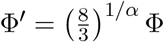, we obtain the power-law form of Eq. 10:

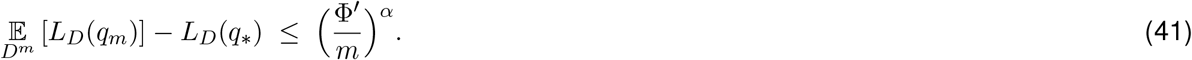

### High-Probability Bound

The oracle regret bound in Eq. 2 can also be presented in a high-probability form, which is more typical in the statistical learning theory literature [46, 55, 28, 29]:

*Given a contextual bandit setup with reward function R*(*s, a*) *and stimulus distribution ρ*(*s*), *consider a Bayesian learner with prior q*_0_(*θ*) *as described above. Then there exist fixed constants* 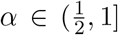 *and c* ≥ 1, *such that for any confidence level δ*∈ (0, 1), *the following inequality holds with probability at least* 1 − *δover samples of S*^*m*^ *from D*^*m*^:

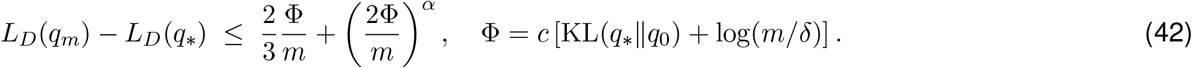

This is a type of Probably Approximately Correct (PAC) Bayes bound, and it can be obtained by applying Markov’s inequality to Eq. 25 before following the same set of steps as in the expectation bound.

### Infimum Oracle Bound

The bound in Eq. 2 compares the learner to the optimal policy distribution *q*_*_. The same argument also yields a comparison with any data-independent policy distribution *q*(*θ*), which need not be optimal. Define the excess regret of any policy distribution *q* and its effective complexity as

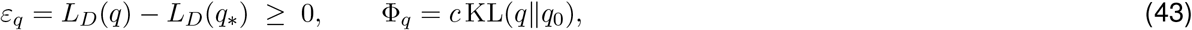

where *ε*_*q*_ ≥ 0 because *q*_*_ achieves the minimal regret. We also have

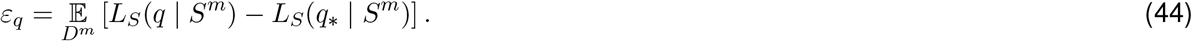

Notice that Eq. 30 holds with *q*_*_ replaced by any *q* on its right-hand side, since the Gibbs posterior minimizes 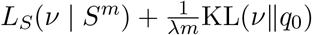 over all *ν*:

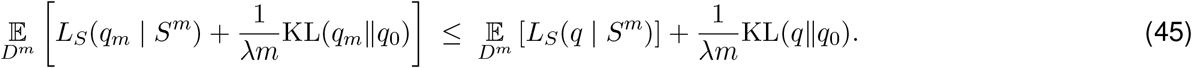

Substituting the above into Eq. 28 and applying the Bernstein assumption and Jensen’s inequality as before, we obtain, in place of Eq. 36,

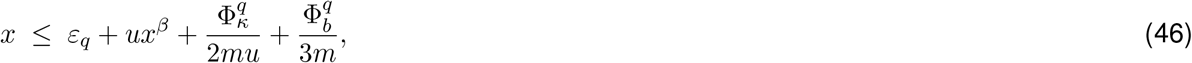

with *x* and *u* defined as before, 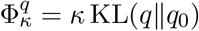, and 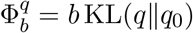. Let 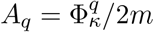 and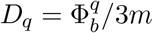. Following the same argument as above, either the first term or the remaining terms dominate the right-hand side, so that

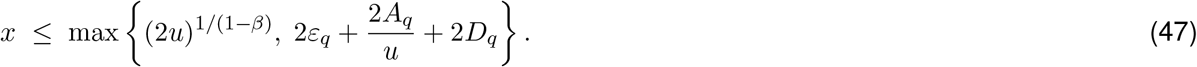

As above, solving the self-bounding inequality gives (with *c* = max(*b, κ*)):

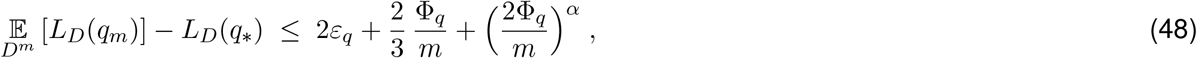

which holds for any *q*. We therefore have:

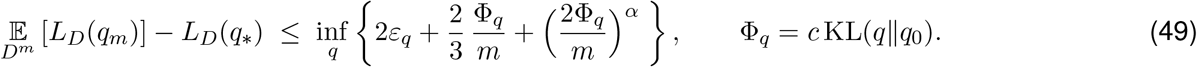

Eq. 49 generalizes Eq. 2, which corresponds to the special case of *q* = *q*_*_, and is at least as tight, because the infimum includes *q*_*_. It offers additional insight into the learning dynamics. The learner’s excess regret Δ(*m*) is bounded by the best trade-off, over all policy distributions, between a distribution’s excess regret *ε*_*q*_ and its effective complexity Φ_*q*_. When *m* is small, the bound is governed by distributions with low effective complexity, even if they are suboptimal; as *m* increases, the best trade-off shifts toward *q*_*_.

### Reward Estimator

The inverse-propensity-score estimator (Eq. 6) is the simplest conditionally unbiased estimator of regret and is sufficient for the theory. In practice, more efficient variations of the estimator that reduce the variance are possible, for example, the doubly robust estimator [61]:

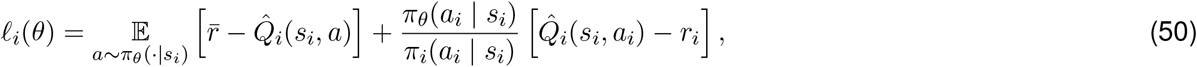

where 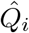 is any estimate of the action-value function. The doubly robust estimator is unbiased if the propensity scores are correct.

Note that the constant *c* = max(*b, κ*) in the bound depends on the regret estimator used by the learner. In the doubly robust estimator, a value model 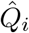 predicts the reward of every action, and the importance-weighted term corrects only the residual error of this prediction on the action actually taken. When the value model is accurate, the residual is small, and so are the range and variance of the estimate, which in turn keeps *c* small.

This is the case in a two-outcome task with deterministic rewards, such as the high-contrast trials of the IBL task. Because the correct choice is always rewarded and the incorrect choice never is, an accurate value model predicts the received reward exactly, 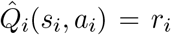, so the correction term vanishes. The estimate then reduces to the probability that policy *θ* would make an error on the current stimulus,

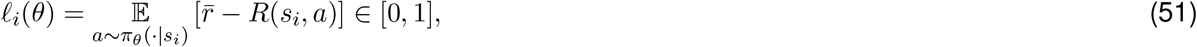

The only remaining source of randomness is which stimulus appears on each trial. As a result, the estimate is bounded, such that *b* ≤ 2. In addition, its variance shrinks in proportion to the policy’s excess regret: since *ℓ*_*i*_(*θ*) ∈ [0, 1] and the optimal regret is zero, the variance is at most E[*ℓ*_*i*_(*θ*)^2^] ≤ E[*ℓ*_*i*_(*θ*)] = *L*_*D*_(*θ*)−*L*_*D*_(*q*_*_). The Bernstein assumption therefore holds with *β* = 1 and *κ* = 1, giving *c* = max(*b, κ*) ≤ 2.

### Continuous Action Space

Lastly, in continuous action (control) settings, *π*_*θ*_(*a* | *s*) is a density over *A*. While the theory is unchanged in this case, in practice, continuous control is often addressed with policy-gradient methods. For example, the REINFORCE algorithm [62] provides an unbiased estimator of the gradient of the expected regret:

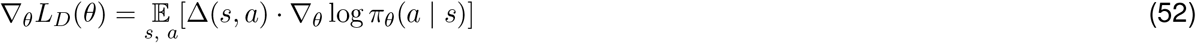

Therefore, a biologically plausible learning rule that descends the gradient of the regret, combined with a regularizer term derived from the prior, can be viewed as approximating the mode (MAP estimate) of the Bayesian posterior rule in Eq. 1. For example:

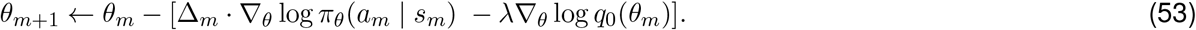

## Supplementary Figures

**Figure S1:**
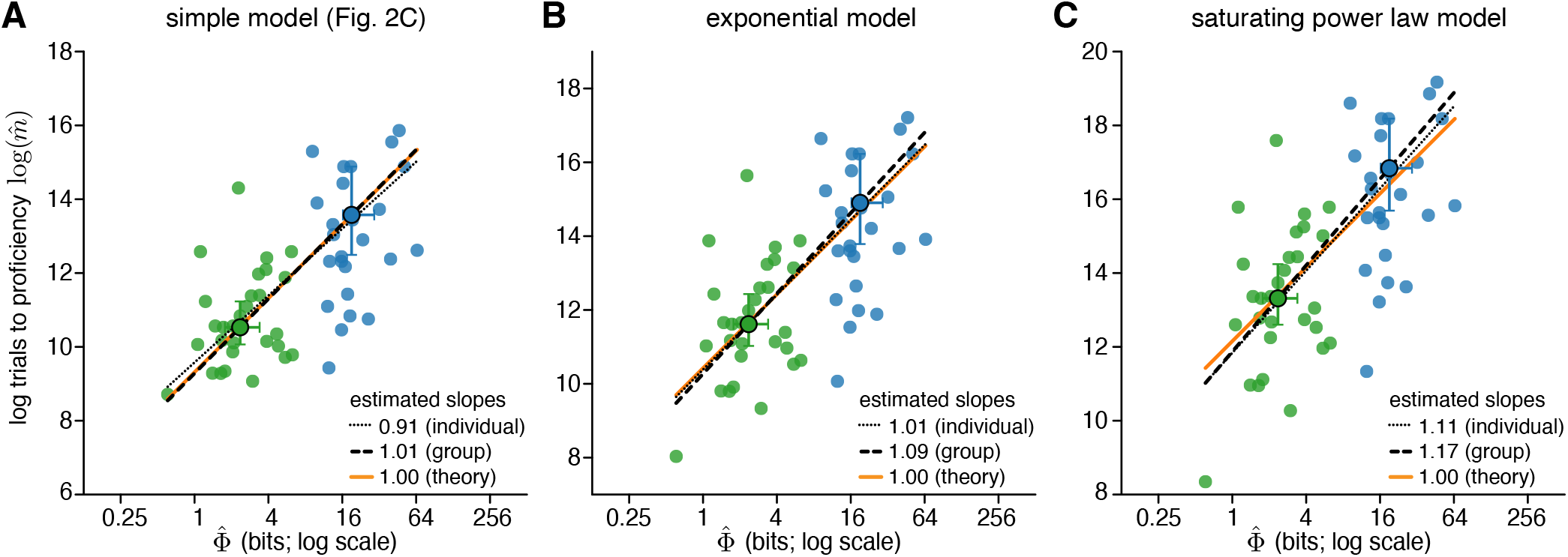
The relationship between measured EC and learning speed is robust to different methods of estimating the number of trials to proficiency. Estimated number of trials to proficiency 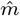 as a function of measured EC 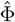, as in Fig. 2C (right), with 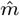 computed under three models of the learning curve (see *Methods*). **A)** Simple model used in the main text,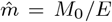. Individual fit slope = 0.91, 95% CI [0.59, 1.22]. **B)** Exponential model. Individual fit slope = 1.01, 95% CI [0.68, 1.14]. **C)** Saturating power-law model. Individual fit slope = 1.11, 95% CI [0.76, 1.47]. In all panels, the orange line shows the theoretical prediction (slope = 1), the black dashed line shows the fit to the task medians, and black dotted line shows the fit across all participants.

**Figure S2:**
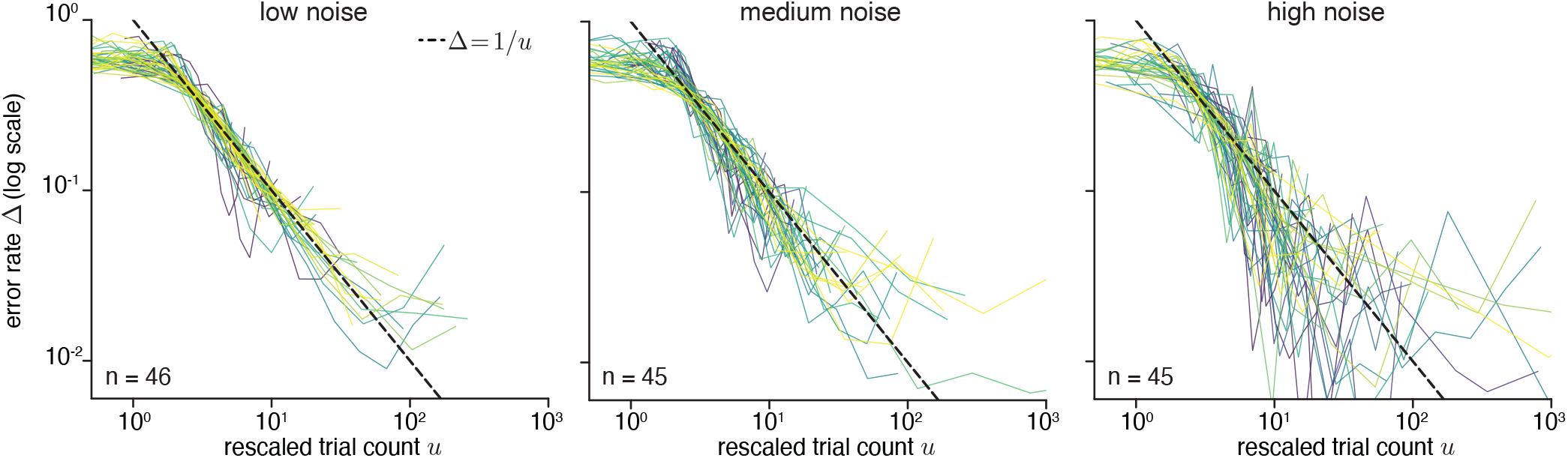
Rescaled learning curves for all mice. Rescaled (*u* = (*m/*Φ^*′*^)^*α*^) learning curves, as in Fig. 3C, for all *n* = 136 mice, shown separately for three groups sorted by the smoothness of their learning curves.

**Figure S3:**
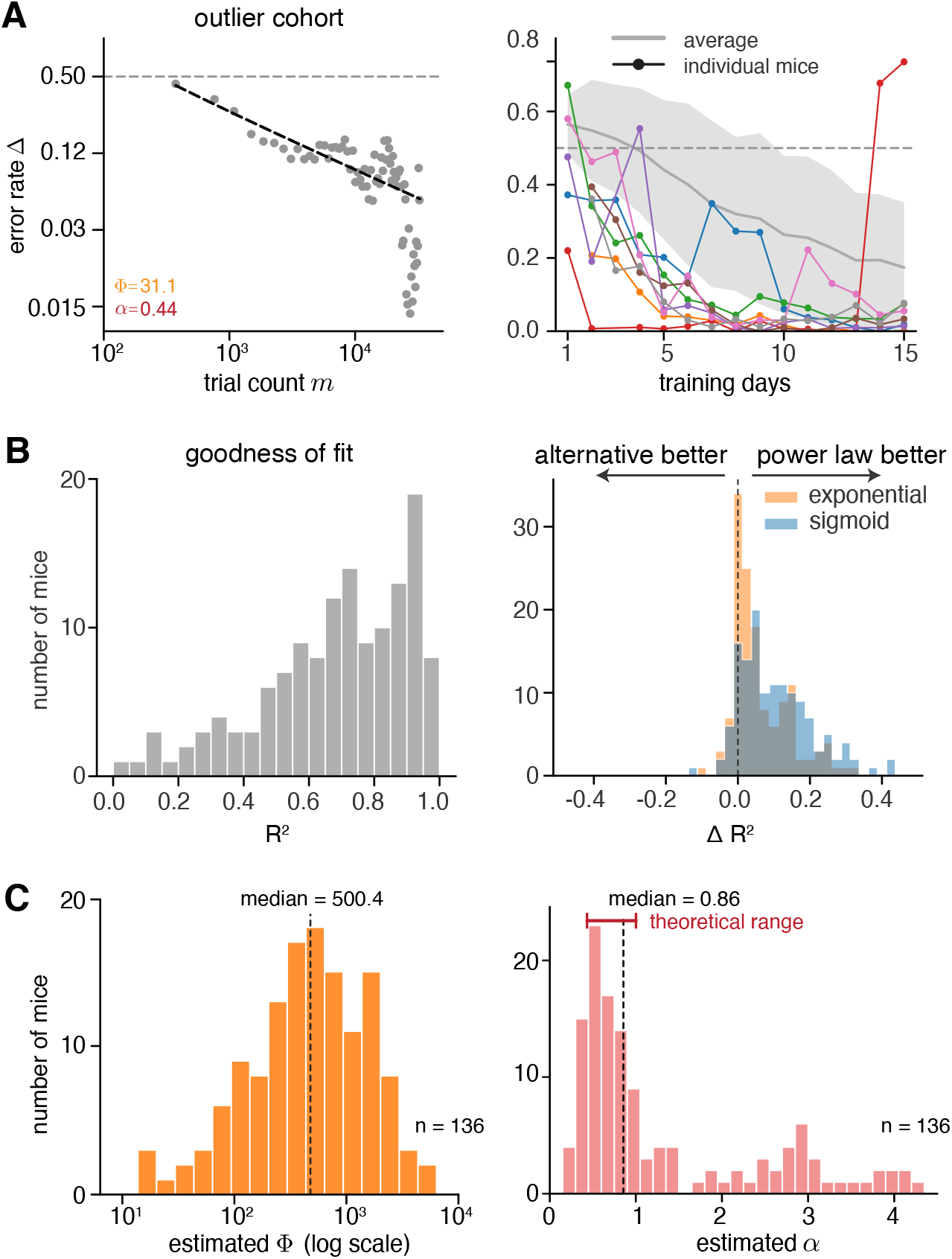
**A)** Average learning curve of the excluded laboratory cohort (left), whose estimates of Φand *α* differed significantly from those of the other cohorts. A likely reason is that many mice in this cohort performed above chance from the early recorded sessions, unlike most mice in the other cohorts (right). Shaded areas indicate ±1 s.d. over all learning curves in the other cohorts. **B)** Left: histogram of the goodness of fit (*R*^2^) for individual mice using the two-term power-law model. Right: difference in *R*^2^ between the power-law model and two other two-parameter learning curve models, the exponential and the sigmoid function (see *Methods*). Positive values indicate a better fit by the power-law model. **C)** Histograms of effective complexity Φand exponent *α* estimated from fits to individual mice (*n* = 136). Dashed lines: medians; red bar: theoretically predicted range of *α* ∈ (1*/*2, 1].

## Notes

https://github.com/lingqiz/inductive-bias

### Competing Interest Statement

The authors have declared no competing interest.

